# *CellDemux:* coherent genetic demultiplexing in single-cell and single-nuclei experiments

**DOI:** 10.1101/2024.01.18.576186

**Authors:** Martijn Zoodsma, Qiuyao Zhan, Saumya Kumar, Javier Botey-Bataller, Wenchao Li, Liang Zhou, Ahmed Alaswad, Zhaoli Liu, Zhenhua Zhang, Bowen Zhang, Cheng-Jian Xu, Yang Li

## Abstract

Multiplexed single-cell experiment designs are superior in terms of reduced batch effects, increased cost-effectiveness, throughput and statistical power. However, current computational strategies using genetics to demultiplex single-cell (sc) libraries are limited when applied to single-nuclei (sn) sequencing data (e.g., snATAC-seq and snMultiome). Here, we present *CellDemux*: a computational framework for genetic demultiplexing within and across data modalities, including single-cell, single-nuclei and paired snMultiome measurements. *CellDemux* uses a consensus approach, leveraging modality-specific tools to robustly identify non-empty oil droplets and singlets, which are subsequently demultiplexed to donors. Notable, *CellDemux* demonstrates good performance in demultiplexing snMultiome data and is generalizable to single modalities, i.e. snATAC-seq and sc/snRNA-seq libraries. We benchmark *CellDemux* on 187 genetically multiplexed libraries from 800 samples (scRNA-seq, snATAC-seq, CITE-seq and snMultiome), confidently identifying and assigning cells to 88% of donors. In paired snMultiome libraries, *CellDemux* achieves consistent demultiplexing across data modalities. Moreover, analysis of 38 snATAC libraries from 149 samples shows that *CellDemux* retains more genetically demultiplexed nuclei for downstream analyses compared to existing methods. In summary, *CellDemux* is a modular and robust framework that deconvolves donors from genetically multiplexed single-cell and single-nuclei RNA/ATAC/Multiome libraries.

## INTRODUCTION

Single-cell sequencing has revolutionized biological research and provided biological insights that would have been indiscernible using bulk-level information: identification of rare cell types, cell-type specific transcriptional programs and chromatin accessibility in health and disease, among others^1^. Droplet-based single-cell and single-nuclei methods are widely used by researchers to encapsulate cells in oil droplets, which are subsequently barcoded to associate sequencing reads to cells^2–11^ to encapsulate cells in oil droplets, which are subsequently barcoded to associate sequencing reads to cells. Droplet-based methods include, but are not limited to, single-cell RNA sequencing (scRNA-seq), single-nuclei RNA sequencing (snRNA-seq) and single-nuclei assay for transposase-accessible chromatin sequencing (snATAC-seq), Cellular Indexing of Transcriptomes and Epitopes by Sequencing (CITE-seq)^12^, as well as the more recent single-nuclei Multiome assay (10x Genomics) that offers paired gene expression and chromatin accessibility data from the same nucleus.

However, single-cell and single-nuclei experiments remain expensive for biological replicates, whereas biological replicates across different conditions are urgently needed to make statistically sound and generalizable conclusions. To address this, an alternative strategy is to multiplex across genetically distinct donors, sequence the composite library and later assign cellular barcodes to the donor of origin based on genetic information. Genetic demultiplexing dramatically reduces unwanted technical variation^13,14^, experimental expenses, and enables inclusion of more biological replicates. Increased numbers of biological replicates or conditions foster more biological diversity and thus generate more generalizable conclusions, which increases statistical power and thus the chances to detect more subtle effects.

Tools to demultiplex pooled single-cell libraries have been developed^13–17^ and widely used by the scientific community^2,3,5–7,9,18–21^. However, currently available genetic demultiplexing tools are designed for scRNA-seq and fail to extend to snRNA-seq or especially snATAC-seq and snMultiome libraries. The extension of demultiplexing tools to snATACseq or Multiome libraries is crucial because of multiple reasons: 1) features between ATAC (peaks) and RNA (genes) libraries differ, 2) ATAC libraries are inherently sparser compared to RNA libraries and require separate processing (e.g. read mapping, genetic variant calling) and 3) current demultiplexing tools do not consider shared barcodes between modalities in snMultiome libraries, which can be utilized to improve demultiplexing performance. Overall, methodological developments on demultiplexing tools are needed to accommodate single-cell and single-nuclei protocols.

In this study, we develop *CellDemux*, a novel, user-friendly computational framework to enable genetic demultiplexing of paired -omics data modalities as well as single-cell and single-nuclei data. This includes pre-processing (e.g., identification of non-empty droplets), confident exclusion of doublets using a consensus approach and assignment of cell barcodes to the donor of origin either within each modality or across modalities. Of particular note, *CellDemux* implements a novel pipeline to demultiplex snATACseq libraries. We show consistent demultiplexing results and validate the results by comparing ATAC demultiplexing results to RNA demultiplexing results in snMultiome libraries, taking advantage of the shared barcodes between the modalities. Overall, we assess the performance and consistency of the proposed framework on scRNAseq, snATACseq, CITEseq and Multiome libraries (combined 7 independent studies, 187 libraries from 800 samples, ∼2M single cells). We show that *CellDemux* can consistently demultiplex donors from single-cell sequencing within and across modalities. Finally, we re-analyze previously published chromatin accessibility data (38 snATAC libraries from three independent studies) to show that this framework outperforms existing methods, giving more reliable single nuclei and statistical power for computational analysis. Our research demonstrates a robust and adaptable framework for genetic demultiplexing that opens the way for more powerful and informative single-cell, single-nuclei and paired -omics experiments. *CellDemux* is freely available at: github.com/CiiM-Bioinformatics-group/CellDemux.

## MATERIAL AND METHODS

### Overview and implementation of the framework

The framework is implemented in Python (v3.9.6) Snakemake (v7.31.0)^22^. We chose Snakemake because of its integration with a wide variety of high-performance clusters and job management systems without changing the underlying code. We containerized most tools to ensure reproducibility using Conda environments, which are freely available. On the user side, input consists of a single excel sheet containing six mandatory entries per library (name, location of the data, number of donors to demultiplex, reference genotype file, data type and optional comment).

We have provided reasonable default setting and computational resources for each tool on Github. Overall, *CellDemux* finished for most libraries within 24 hours, starting from cell calling to matching to the reference genotype. We have made this workflow modular, meaning some steps can be skipped depending on the scenario to alleviate the computational burden.

### Data pre-processing

Pre-processing of the published datasets is described in their respective publications^2,4,21^. For the unpublished datasets we used CellRanger (v7.1.0), CellRanger-atac (2.1.0) and CellRanger-arc (v2.0.2) with default parameters for RNA, ATAC and Multiome libraries respectively.

### Methods to estimate ambient RNA contamination

#### Soupx

We used SoupX^31^ v1.6.2 with default parameters, except forceAccept = T, soupQuantile = 0.1 and tfidfMin = 0.0 to avoid errors when encountering libraries with high contamination or where too few markers were found to estimate contamination. SoupX was only used on RNA libraries, for Multiome / CITEseq libraries we first subsetted the matrix that was passed to SoupX. Over all 149 RNA libraries, we observed that SoupX estimates of contamination were highly correlated to Souporcell estimates (Pearson’s r: 0.83, p < 2.2 x 10^−16^)

### Methods to identify non-empty droplets

#### CellBender

We ran CellBender^24^ within a Singularity container on GPUs to increase its efficiency. We ran CellBender only on RNA libraries and subsetted Multiome/CITEseq libraries prior to running CellBender. CellBender was run with arguments: --expected-cells 8000, --cuda, --cells-posterior-reg-calc 10, --posterior-batch-size 2, --epochs 150. We found that CellBender is not particularly sensitive to these parameters, and 150 epochs were always enough to reach convergence.

#### EmptyDrops

EmptyDrops^23^ (R package DropletUtils v1.18.1) was run on RNA libraries to estimate non-empty oil droplets. Prior to using EmptyDrops we subsetted CITEseq or Multiome libraries to only include RNA counts. We used a False Discovery Rate (FDR) of 0.001 for cell calling as recommended by the EmptyDrops authors. Other arguments were left as default.

For ATAC libraries, we used the *CellRanger* ATAC cell calling algorithm. We attempted to use RNA-based tools *CellBender* and *EmptyDrops* to ATAC sequencing results, and observed that both tools identified mostly cell-containing droplets unique to data modalities in Multiome libraries (Supplemental Figure 1B). Meaning that, while the cell-containing droplets should be largely overlapping, RNA-based tools were not able to identify the correct droplets in ATAC. Hence, we continued with the CellRanger ATAC cell calling results.

### Methods to identify doublets

#### Demuxlet

We used the Popscle suite (https://github.com/statgen/popscle) to run Demuxlet. We used several helper tools for Popscle in pre-processing the genetic data created by the Aerts lab (at: https://github.com/aertslab/popscle_helper_tools). Briefly, we filter out from the genotype reference anything that is not a single nucleotide polymorphism (only_keep_snps function), filter mutations that do not vary across all samples (filter_out_mutations_homozygous_reference_in_all_samples, filter_out_mutations_homozygous_in_all_samples) and calculate the allele frequencies, allele counts and allele numbers (calculate_AF_AC_AN_values_based_on_genotype_info). This leaves a set of informative variants with discriminative power between samples. Next, the bam file is filtered for the appropriate barcodes using popscle_filterbam.sh (popscle). We produced the pileup using popscle dsc-pileup and subsequently demultiplexed using popscle demuxlet.

#### Vireo

Vireo^15^ and cellsnp-lite v1.2.3 (htslib version: 1.17) was ran using default parameters and according to the author’s instructions.

#### Amulet

We ran Amulet^27^ v1.1 using the shell script provided by the authors with default parameters. The human autosomes and blacklist regions provided to Amulet were also supplied by the authors and are available on Github (https://github.com/UcarLab/AMULET).

#### ArchR

ArchR^28^ v1.0.2 was used to estimate doublets in ATAC libraries. We first created Arrow files using the fragment files for each ATAC library using default parameters. The valid barcodes we supplied to ArchR were the non-empty droplets as estimated by CellRanger. Then, we estimated doublet scores per nucleus (addDoubletScores function) using k=10, knnMethod = UMAP and LSImethod = 1, the default values. Nuclei were removed at the lenient threshold of DoubletEnrich score > 1.

#### Souporcell

We ran the Souporcell^13^ pipeline manually to estimate singlet following the author’s instructions. For different libraries, we map the reads using mappers suited to each data modality: Minimap2^32^ (arguments: -ax splice -t 15 -G50k -k 21 -w 11 –sr -A2 -B8 -O12,32 -E2,1 -r200 -p.5 -N20 -f1000,5000 -n2 -m20 -s40 -g2000 -2K50m –secondary=no) for RNA and BWA MEM^33^ (arguments: -t 15) for ATAC. From these remapped reads, variants were identified using Freebayes^34^ (see below). Vartrix was used to count the alleles using the identified variants with default parameters. Finally, cells were clustered using the Souporcell cell clustering algorithm to identify singlets.

### Methods to call variants

We used FreeBayes^34^ for all results presented. Freebayes was run with arguments: -iXu -C 2 -q 30 -n 3 -E 1 -m 30 –min-coverage 6 on bam files resulting from RNA-seq and ATAC-seq alignments. Variant calling was done per chromosome in parallel using freebayes-parallel.

Moreover, we tested a recently published algorithm, Monopogen^35^, to call variants in single-cell and single-nuclei sequencing data. Consistent with the original publication, we found that Monopogen called more variants in ATAC-seq data compared to RNA-seq data. Compared to FreeBayes, Monopogen identified similar number of variants in RNA-seq data but higher number of variants per library in ATAC-seq data (Supplementary Figure 4).

Both variant callers are included in CellDemux and can be interchangeably used.

### Cell clustering

Having identified a confident set of singlets for each ATAC and RNA library, we manually ran Souporcell again manually to finally cluster cells/nuclei. We filter the mapped reads (RNA: Minimap2^32^, ATAC: BWA MEM^33^) for the appropriate barcodes and run Freebayes^34^ using the same parameters specified above. Vartrix and Souporcell’s cell clustering algorithm were re-run using default parameters to identify the final cell clusters. Of note, we do not consider the ambient RNA estimations from this Souporcell output since the input sequence data was filtered and will therefore skew estimations.

### Matching cell clusters to donors

To assign the cell clusters to donors, we match the genotypes of the cell clusters to reference genotypes. First, we identify a set of variants that are present in both genotypes based on chromosome, chromosomal position, alternate allele and reference allele. Both genotype files are filtered for this set of variants. Following, we systematically compare the genotypes for this set of variants. Because we cannot distinguish between the paternal and maternal alleles, we collapse the genotypes to the count of alternate alleles per variant in both the inferred and reference genotypes. We sum the count of alternate alleles over these variants for every combination between cell clusters and donors. To statistically identify outliers (e.g., a donor that matches more variants to a cell cluster compared to other donors), we use Grubbs’ test^36^. We test only the donor with the highest number of variants per cluster. Unadjusted p-values < 0.05 were considered significant outliers and therefore indicated matching of a cell cluster to a donor. We manually checked whether the assigned donor agreed with the single-cell experimental design.

Prior to matching the cell cluster genotypes to the reference genotype, we recommend imputing the reference genotypes to maximize the number of variants to consider. We observed that imputing the reference genotypes dramatically increased the number of matching variants for all donors and more confidently showed the matching donor in that library.

## RESULTS

### A comprehensive and user-friendly framework to enable genetic demultiplexing

We developed *CellDemux*, a user-friendly and comprehensive computational framework to enable assignment of cells to genetically different donors from single-cell, single-nuclei and paired -omics libraries with mixed donors (Figure 1). The framework, implemented in the workflow management system Snakemake^22^, supports a wide range of data for genetic demultiplexing, including scRNA-seq, snATAC-seq, CITE-seq and paired snMultiome data. Starting from raw sequencing data, *CellDemux* identifies cell-associated droplets by discarding droplets contaminated by ambient RNA. Contamination of cell-free ambient mRNA molecules is a challenge in single-cell experiment, mainly because the real expression profile is masked which confounds further downstream analyses. Particularly in the case of genetic demultiplexing, ambient RNA interferes with the estimation of genotypes from single-cell data as a particular genotype may be contaminated with false-positive reads derived from ambient RNA. Therefore, *CellDemux* implements two methods (*EmptyDrops* and *CellBender*) to confidently separate empty vs non-empty droplets. Next, cell barcodes containing a single cell or nucleus are identified using a consensus approach of computational tools specifically designed for each data modality. This is important because heterogenic doublets, those including cells from different donors, interfere with the genotypic estimations in that cell/nucleus as the sequenced reads harbor different alleles for a genetic variant. Therefore, *CellDemux* utilizes different doublet callers to identify and remove the cell barcodes that likely contain more than one cell/nucleus in order to retain a set of high-confidence singlets. Finally, we cluster these singlets on genotypic information^13^ and match the inferred and reference genotypes, assigning barcodes to donors across data modalities. We compare demultiplexing results across modalities to show the consistency of cell clustering and donor assignments between modalities.

**Figure 1.**
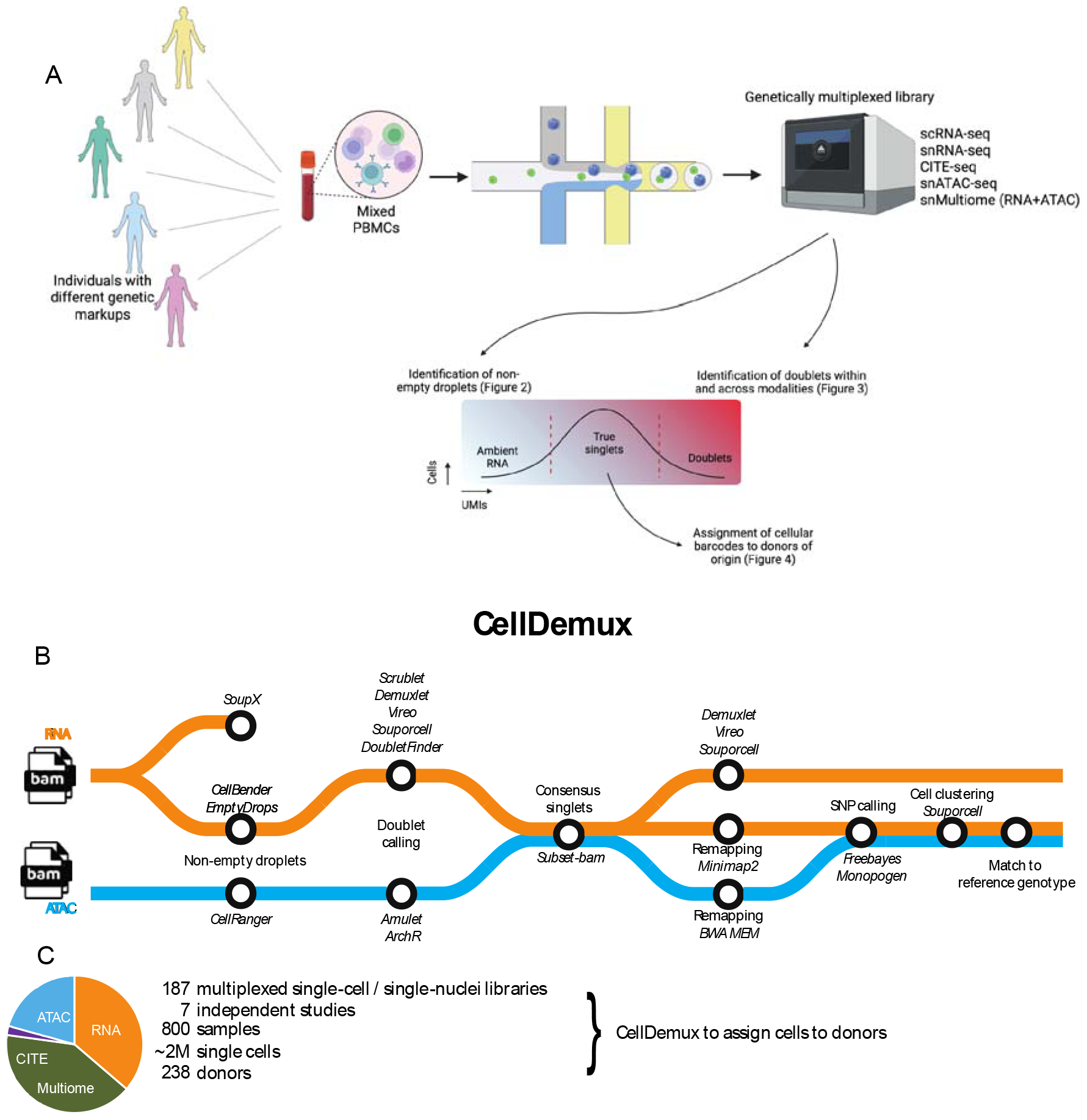
Study design. (A) Schematic overview of the study workflow and aims. Genetically varying individuals are mixed and sequenced (scRNA-seq, snRNA-seq, snATAC-seq, snMultiome, CITE-seq) and later demultiplexed to assign cells back to the donor of origin. Droplets that contain only ambient RNA are removed, as are droplets that likely contain multiple cells. High-quality singlets are subsequently used for further demultiplexing. (B) Overview of CellDemux, a computational framework to confidently assign cell barcodes from genetically multiplexed single-cell or single-nuclei experiments to their respective donor. (C) Data used in the current study includes 189 genetically multiplexed libraries, covering scRNAseq, snATACseq, CITEseq and snMultiome libraries.

We benchmark *CellDemux* on 187 multiplexed libraries of single-cell and single-nucleus sequencing data from 7 studies, covering 800 samples from 238 genetically different donors and approximately 2M single cells (both published^2,4,21^ and unpublished data, Supplemental Table 1). Data from these studies include scRNA-seq (66 libraries), CITE-seq (4 libraries), snATAC-seq (38 libraries) and paired RNA+ATAC snMultiome (79 libraries).

### Confident identification of non-empty oil droplets within and across modalities

Identification of high-quality cells/nuclei is crucial in genetic demultiplexing to identify the donors mixed in a sequencing library. The first computational step in droplet-based single-cell sequencing is identifying the oil droplets that likely contain cells. By comparing the output of different cell calling tools across modalities we aim to assess their consistency and identify high-quality cell-associated droplets.

For 149 RNA libraries (either scRNA or the RNA part of a snMultiome/CITE-seq library), *CellDemux* implements two widely employed tools to estimate non-empty droplets: *EmptyDrops*^23^, used by *CellRanger* (10X Genomics), estimates deviations from the ambient RNA pool to identify empty droplets, while *CellBender*^24^ is a deep generative model that learns the background noise profile. *EmptyDrops* overestimated the number of non-empty droplets (up to ∼80K non-empty droplets per library) especially in pools with higher estimated ambient RNA contamination (Figure 2B, Supplemental Figure 1A). Overall, there was moderate overlap in the identified non-empty droplets between *EmptyDrops* and *Cellbender* (47.5% consistency, Figure 2A). Results from both tools diverged more as the estimated ambient RNA contamination increased (Figure 2C). To identify non-empty droplets in 119 snATAC libraries (either snATAC or the ATAC part of a snMultiome library), we used the ATAC cell calling algorithm from *CellRanger*. Finally, we applied *CellDemux* to snMultiome libraries with shared barcodes between RNA and ATAC to assess the consistency between RNA/ATAC non-empty droplets. Non-empty RNA droplets called by *CellBender* overlapped significantly more with the ATAC non-empty droplets compared to *EmptyDrops* (Wilcoxon ranked-sum test, p = 0.011) (Figure 2D).

**Figure 2.**
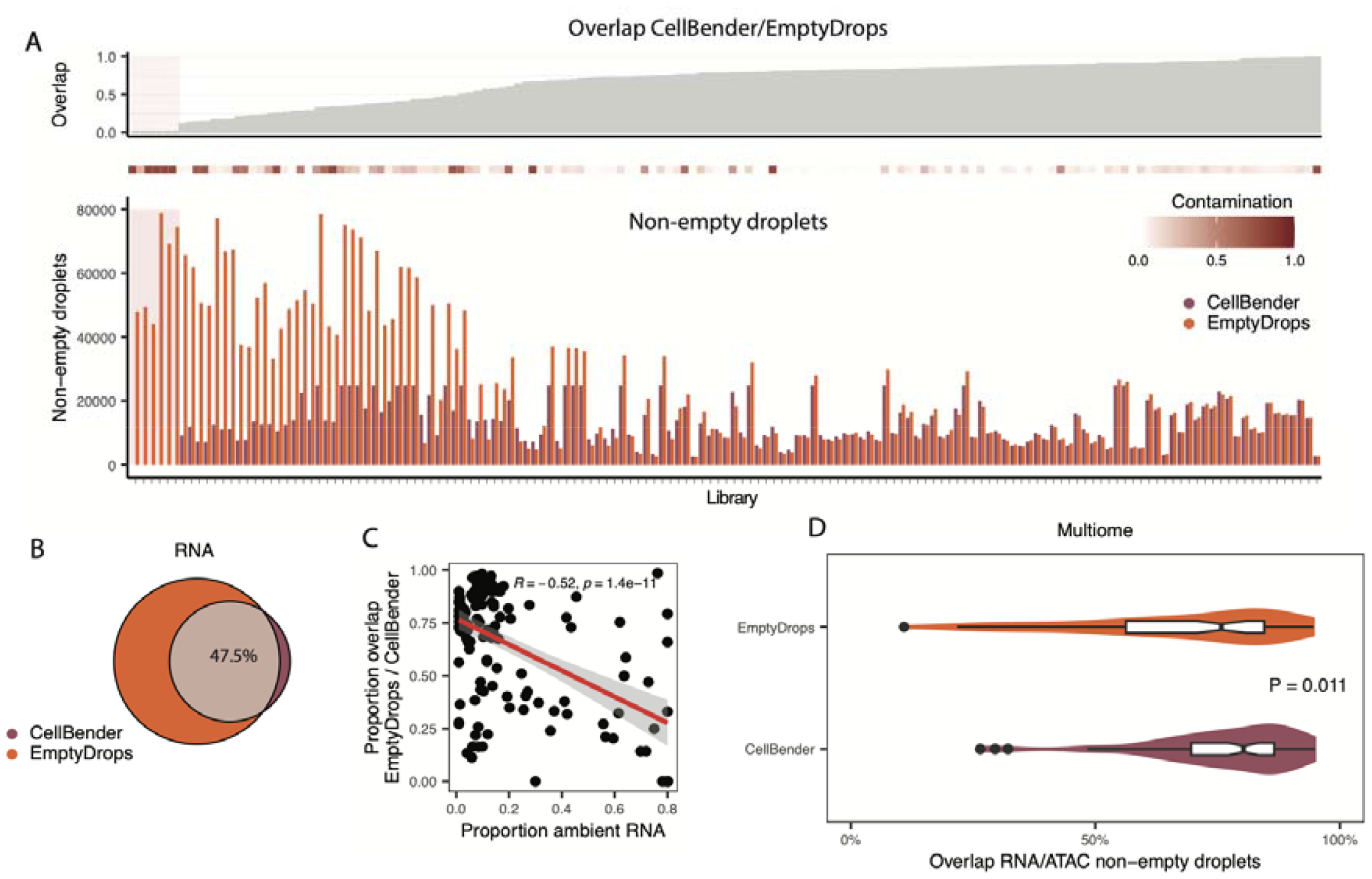
Identification of non-empty droplets across data modalities. (A) Overlapping in non-empty barcodes between CellBender and EmptyDrops across 149 RNA libraries (either RNA or as part of a Multiome experiment). (B) For each of the 149 RNA libraries, the number of non-empty droplets estimated by CellBender and EmptyDrops (bottom), the estimated ambient RNA contamination (middle) and the overlap between CellBender/EmptyDrops (top). Libraries shaded in red indicate the CellBender failed to converge and did not identify any cells. No further analyses were performed on these 5 libraries (far left side) (C) Correlation between the estimated ambient RNA contamination per library and the overlap between CellBender/EmptyDrops estimations of non-empty droplets. Each dot represents one of 149 RNA libraries. (D) Overlap between RNA and ATAC barcodes across 79 Multiome libraries. RNA barcodes were identified either by EmptyDrops or CellBender, ATAC barcodes were identified using CellRanger’s standard cell calling pipeline. Center line in each boxplot is the median, bounds are the 25th and 75th percentiles (interquartile range).

Overall, these results suggest substantial variabilities in the identification of non-empty droplets dependent on data quality. *CellDemux* implements different cell calling methods suited to each data modality, and therefore provides high-quality non-empty droplets per library. We consider for further analysis a set of confidently called non-empty droplets (RNA libraries: CellBender ⋃ EmptyDrops, ATAC libraries: CellRanger).

### Removal of doublets within and across modalities

To further ensure high quality of the sequencing data prior to demultiplexing, *CellDemux* employs a consensus approach using well-established doublet calling tools suited for each modality to identify and remove doublets in single-cell and single-nuclei data. Specifically, *CellDemux* implements : *Vireo*^15^, *Demuxlet*^14^, *Souporcell*^13^, *DoubletFinder*^25^, *Scrublet*^26^ for RNA libraries, and *Amulet*^27^ and *ArchR*^28^ for ATAC libraries. We compared the outcome of these tools within and across modalities to identify a confident set of singlets for further demultiplexing.

For 70 RNA/CITE-seq libraries, there was notable variation in terms of the total number of singlets, with a maximum 1.7-fold-difference (260K single cells) across tools, and the extent of overlapping singlets between the different methods (Figure 3A). In ATAC libraries, *Amulet* identified ∼110K singlets more on the same datasets compared to *ArchR*’s doublet calling method. The large majority of singlets identified by ArchR were shared with *Amulet* (Figure 3B). These variable results may come from varying statistical power to detect doublets, and underlines that a consensus approach is necessary when identifying doublets. Relying solely on the performance of a single tool will likely lead to both false positive and false negative doublet calls.

**Figure 3.**
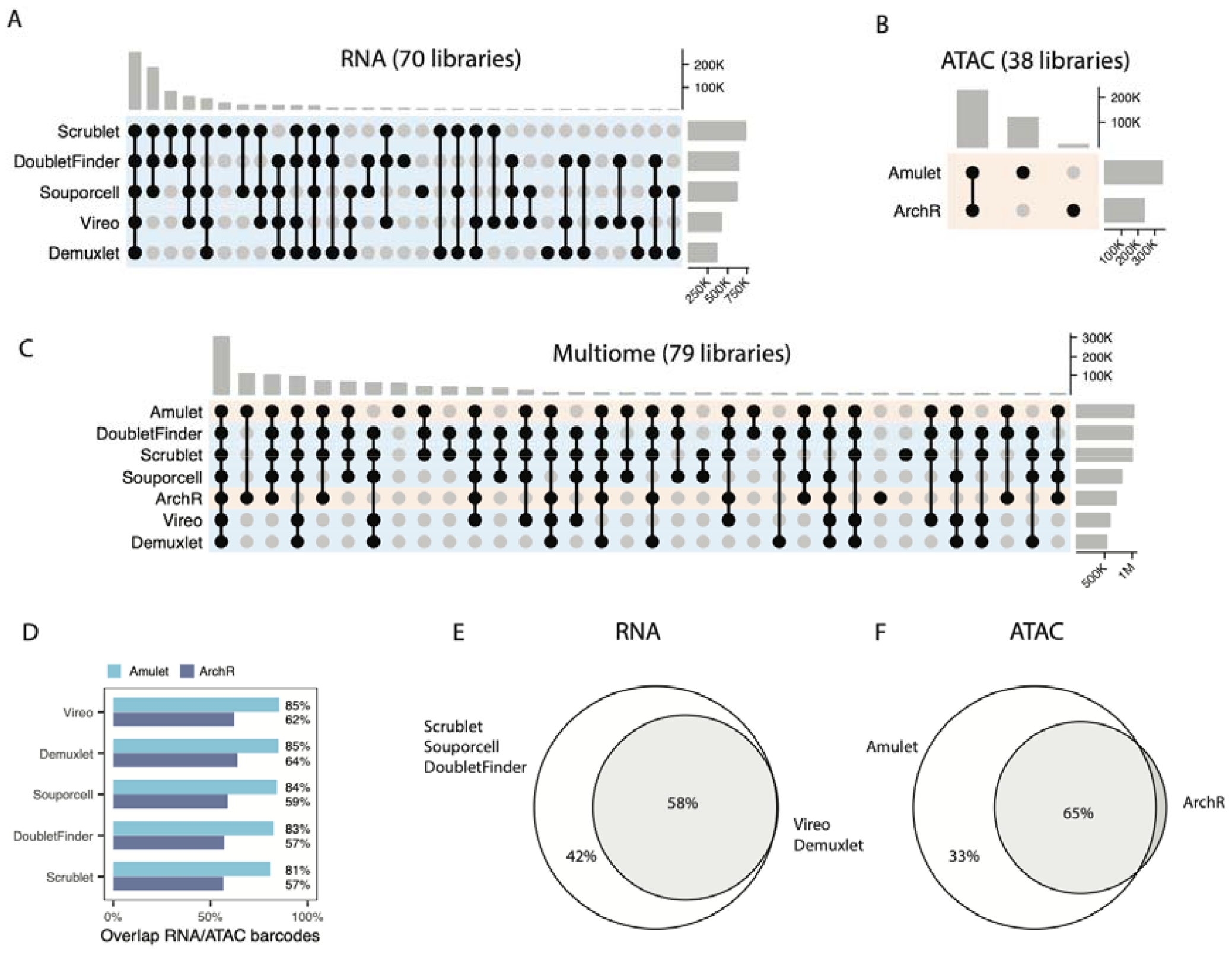
Confident identification of singlets across data modalities using a consensus approach. UpSet plots depicting doublet calls in each data modality across 70 RNA libraries (either scRNA-seq or part of CITE-seq/snMultiome libraries) (A), 40 ATAC libraries (either snATAC-seq or part of snMultiome libraries) (B) and 79 RNA+ATAC snMultiome libraries (C). The top bars indicate the total number of singlets in this intersection, the side bar plots indicate the total number of singlets identified per tool. (D) Barplot showing the concordance between RNA and ATAC for each of the doublet calling tools, as the proportion of identified singlet barcodes by RNA that are also singlets in ATAC by either of the tools. (E) Overlap between singlets identified by Souporcell/Scrublet/DoubletFinder versus Vireo/Demuxlet. (F) Venn diagram showing the overlap between identified singlets by Amulet or ArchR.

We again took advantage of the shared barcodes between RNA and ATAC in snMultiome libraries to assess the performance of *CellDemux* based on the concordance of doublet predictions across modalities. Overall, a large subset of singlets was independently identified across modalities and different doublet calling tools, suggesting that these are high-confidence singlets (Figure 3C). Remaining sets of singlets varied considerably and were classified as singlets in only one modality or by a subset of doublet callers. For example, snRNA-seq singlets were more concordant with Amulet singlets compared to ArchR across all doublet calling tools, but we did not observe differences in proportional overlap between each of the RNA doublet callers (Figure 3D).

We noticed that singlets identified in RNA libraries by *Demuxlet/Vireo* were also identified by *Scrublet, Souporcell* and/or *DoubletFinder*, which identified an additional 322K singlets (Figure 3E). Similar for ATAC libraries, >98% of the singlets identified by *ArchR* were also identified by *Amulet*, who identified up to 116K single nuclei extra (Figure 3F). Furthermore, assessing the paired snMultiome libraries shows that these singlets are largely shared across modalities and identified by several independent methods (Supplementary Figure 2C).

Based on these cross-modal validation results, we therefore consider for further genetic demultiplexing a broad set of singlets (RNA: Souporcell ⋃ Scrublet ⋃ DoubletFinder, ATAC: *Amulet*, Multiome: Souporcell ⋃ Scrublet ⋃ DoubletFinder ⋃ *Amulet*). We propose a broad set of singlets to not exclude potential real singlets (i.e. false positive doublets), who will be filtered out later based on genotypic information.

### Assignment of single cells and nuclei to donors in genetically multiplexed libraries across modalities

Having established a confident set of singlets per library by identifying non-empty droplets (Figure 2) and singlets (Figure 3), we cluster cells based on genetic variation and use reference genotypic information to assign cellular clusters to donors. We chose the Souporcell^13^ model to cluster cells because of its flexibility (quality control, processing, mapping and variant identification can be tailored to data modalities as we have demonstrated here) and previously demonstrated superior demultiplexing results on single-cell RNA sequencing data^13^. Per cluster, we call variants and systematically compare the inferred and reference genotypes to count the matching number of variants per cluster and donor (Methods). Finally, we manually verified that the demultiplexed donors were included in that library.

Application of *CellDemux* to 187 genetically multiplexed libraries from seven independent studies shows that we significantly match 92% of cell clusters to donors in 70 RNA/CITE-seq libraries (Supplemental Figure 3A), 95% in 38 ATAC libraries (Supplemental Figure 3B) and 84%/93% (RNA/ATAC) in 79 paired RNA+ATAC snMultiome libraries (Figure 4A). We observed few libraries (ATAC: 1, RNA: 1, Multiome: 2) with inconclusive results, where 1) multiple cell clusters matched to the same donor, suggesting poor cell clustering or samples with poor viability, or 2) two samples matched almost equal number of variants to cell clusters, suggesting a possible sample swap in the genotyping or poor clustering of cells. Overall, the number of nuclei recovered and assigned to donors was consistent between RNA and ATAC in the snMultiome libraries. Donors that were not demultiplexed from the single-cell data were often not demultiplexed in multiple modalities, suggesting poor sample quality rather than demultiplexing artifacts. Aside from these factors, there remain libraries where not all donors were identified or lesser number of cells were assigned to donors. This is possibly due to the fact that these libraries were characterized by higher estimated ambient RNA contamination, lower mapping quality of the sequenced reads and lower number of SNPs identified in the single-cell data (Figure 4A).

**Figure 4.**
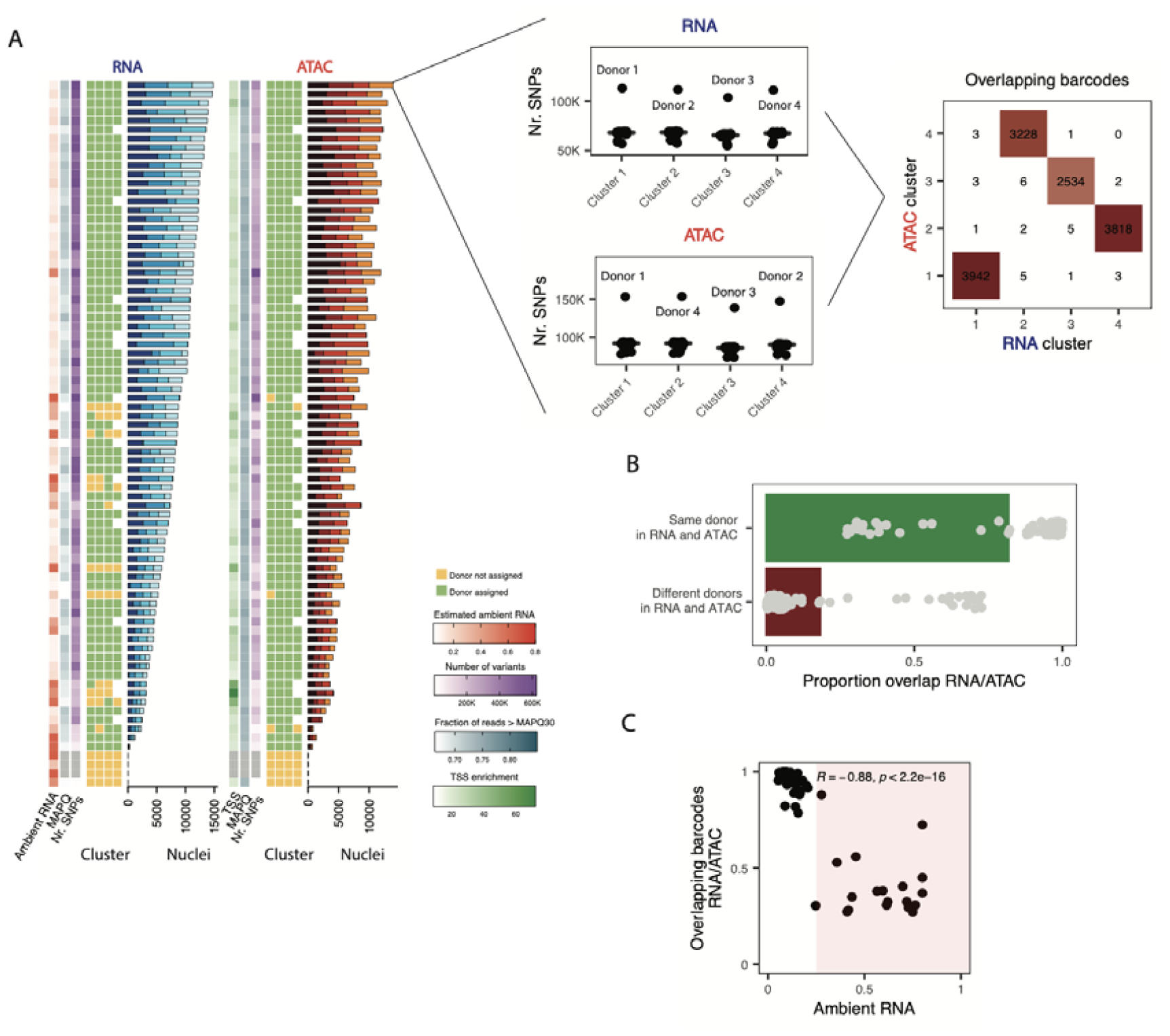
Assigning cell barcodes to donors in multiplexed libraries. (A) Composite heatmap showing demultiplexing results across 79 RNA+ATAC snMultiome libraries. Each row represents a paired RNA (left) and ATAC (right) snMultiome library, where the heatmap shows whether a cluster of cells was significantly matched to a donor based on genotypic information (green) or not (yellow). Furthermore, we annotate each library with the estimated ambient RNA contamination (RNA only), transcription start site enrichment (ATAC only) as well as the fraction of reads that are > MAPQ 30 and the number of variants identified in the sequencing data by FreeBayes. Barplot on the right shows the number of nuclei assigned to each donor in that library. We highlight one library and show how the genetic matching assigns donors to clusters (Methods), and how barcodes overlap between donors across modalities. (B) Fraction of barcodes that are consistently assigned to the same donor in RNA and ATAC independently. Each dot represents one of 79 Multiome libraries, bars indicate means. (C) Dot plot showing the correlation between the estimated ambient RNA contamination and the fraction overlapping reads between RNA and ATAC (e.g., fraction of reads assigned to the same donor, independently across two modalities).

Comparing across modalities in the paired snMultiome libraries showed that cell clustering and matching cells to donors was consistent between RNA and ATAC, where on average 84% of barcodes were assigned to the same donors in both modalities (Figure 4B). This shows that *CellDemux* is able to consistently deconvolve donors in multiplexed libraries across modalities. We further assessed the confounding factors that could influence the consistency between modalities, and found that libraries with lower agreement between RNA and ATAC were more contaminated with ambient RNA, suggesting lower quality of that specific library (Pearson’s r: -0.88, p < 2.2e-16) (Figure 4C).

Assignment of barcodes to donors in RNA libraries was consistent with results obtained from Vireo and Demuxlet (Supplementary Figure 2A), but *CellDemux*, using the Souporcell model, was able to assign more cells to donors compared to either tool (Supplementary Figure 2B).

### CellDemux outperforms existing methods on snATAC-seq data

One of the unique features of *CellDemux* as a demultiplexing tool is its applicability to different modalities of single-cell and single-nuclei data. Single-cell RNA demultiplexing tools are widely used^13–15,29^ but do not extend to snATAC-seq data. To show that *CellDemux* outperforms existing methods on snATAC-seq data, we compare results to three snATAC-seq datasets previously generated, analyzed (standard Souporcell pipeline, the best that was available at time of publishing) and published by our lab^2,4,21^. Overall, these three studies encompass 32 snATAC-seq libraries that are multiplexed across genetically distinct donors (163 samples).

*CellDemux* is able to demultiplex more samples from snATAC-seq data compared to existing tools. We compared the number of samples that were significantly demultiplexed for three independent studies where samples from 3-6 donors were pooled for snATAC-seq. *CellDemux* identified and assigned cells to almost all samples per library across all three studies, i.e., study 1: 20/22 (91%) samples, study 2: 75/80 (94%) samples and study 3: 47/47 (100%) of samples demultiplexed. This is in contrast to the previously published results based on the standard Souporcell pipeline that identified fewer samples in two out of the three studies (study 1: 16/22 (73%), study 2: 49/80 (61%) and study 3: 47/47 (100%) samples multiplexed) (Figure 5A). This could be due to lower sequence mapping quality, lower number of SNPs identified and subsequent poorer clustering of cells using the standard Souporcell pipeline on snATAC-seq data, compared to the improved snATAC-seq-specific pipeline used by *CellDemux*.

**Figure 5.**
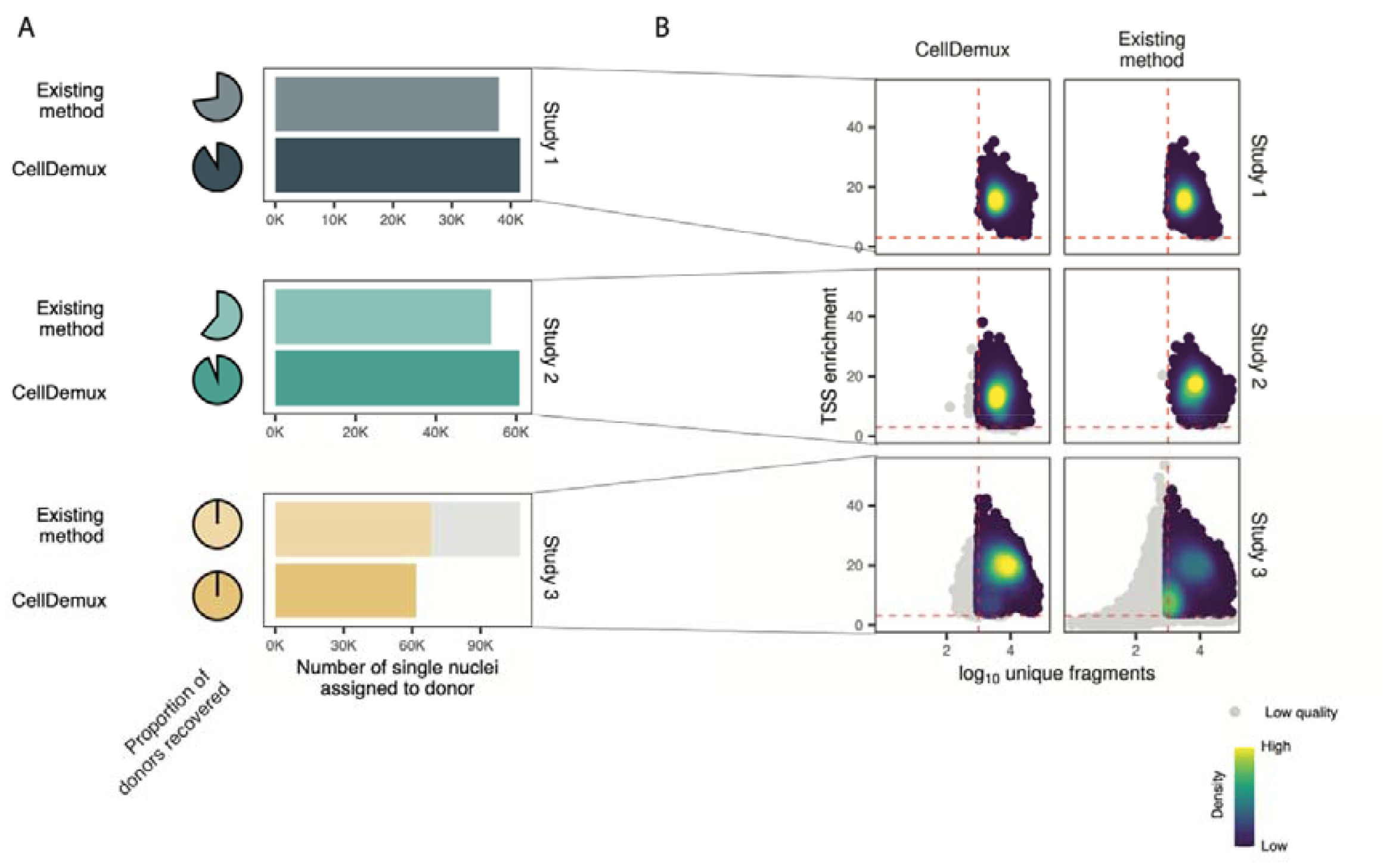
Re-analysis of previously published data shows that CellDemux outperforms existing methods. (A) Proportion of donors recovered (pie charts) and number of singlets assigned to donors per study, for CellDemux and the previously published using the standard Souporcell pipeline on the same snATACseq libraries for three independent studies. We considered only high-quality singlets. Low-quality singlets that are filtered out in downstream steps are colored gray (TSS enrichment < 4, or unique number of fragments per cell < 1000) (B) Dot plot showing the number of unique fragments per nucleus versus the transcription start site enrichment (TSS) for each of the three studies (rows), colored for density. We considered singlets assigned to donors based on CellDemux or using the previously published standard Souporcell pipeline results. Cells that are low quality (TSS enrichment < 4 or number of unique fragments per cell < 1000, red lines) are colored gray.

CellDemux outperforms existing tools in accurately identifying high-quality nuclei from snATAC-seq data. We compared the number of single nuclei that are retained for downstream analyses between *CellDemux* and the previously used standard Souporcell pipeline (Figure 5B). *CellDemux* identified more singlets in two out of the three studies (study 1: 3599 (8.6%) single nuclei, study 2: 7031 (10%) single nuclei more assigned to donors). In contrast to studies 1 and 2, *CellDemux* assigned fewer nuclei to donors in study 3 compared to previously published results. A more in-depth assessment of the demultiplexed single nuclei showed that *Souporcell* previously overestimated the number of singlets (∼30.000 singlets in a single library). These singlets were low-quality as defined by the number of unique fragments and the TSS enrichment per nuclei, and are later filtered out in downstream analyses. It is likely that there are more low-quality singlets among the demultiplexed singlets by *Souporcell* that we do not filter out based on only TSS enrichment and the number of unique fragments.

In summary, we showed here that *CellDemux* enables genetic demultiplexing of single-nuclei chromatin accessibility data. By leveraging methods specific to ATAC-seq, CellDemux substantially improved demultiplexing results compared to existing methods, both in terms of nuclei retention and accurately demultiplexed donors.

## DISCUSSION

In this study, we presented the *CellDemux* framework that enables confident demultiplexing of single cells and nuclei to genetically different donors across data modalities. *CellDemux i*s implemented in the workflow management system Snakemake to facilitate integration with high-performance clusters and job management systems. Furthermore, each module is self-contained to ensure maximal reproducibility.

We have evaluated *CellDemux* across a wide range of genetically multiplexed libraries including scRNA-seq, CITE-seq, snATAC-seq and paired RNA+ATAC snMultiome data. These libraries comprised a set of challenging scenarios (e.g., high ambient RNA contamination, variability in sequencing quality, varying number of donors, different data modalities) and therefore present robust test cases. We showed that *CellDemux* consistently demultiplexes donors from genetically multiplexed single-cell and single-nuclei sequencing. Assessments of paired snMultiome libraries showed that *CellDemux* assigns highly concordant sets of barcodes to donors across modalities. Finally, re-analysis of previously demultiplexed ATAC libraries showed that *CellDemux* confidently demultiplexes donors and recovered more high-quality nuclei that would have otherwise been discarded.

Genetically multiplexed single-cell and single-nuclei libraries comprise several types of doublets: homotypic and heterotypic based on cell type, as well as homogenic and heterogenic based on genotype. Different doublet callers have varying detection power for each class of doublets. For example, Souporcell^13^ was developed for genetically multiplexed libraries and explicitly models the allelic fractions between clusters to identify heterogenic doublets. Homogenic doublets could be missed using this approach but can be identified by other tools such as *Scrublet*^26^ or *DoubletFinder*^25^. The idea of using multiple bioinformatic tools to identify doublets in droplet-based single-cell sequencing has been suggested before and was shown to improve consistency and results^29,30^. However, previous research remained limited to single-cell RNA sequencing^29^ and hashtag-based demultiplexing methods^30^ and did not include other library types such as ATAC-seq or snMultiome. We considered in this work a broad set of singlets per library by considering multiple modality-specific tools. Consequently, our results become less dependent on the outcome of a single tool and therefore more robust.

While there is a plethora of doublet-calling tools for single-cell or single-nuclei RNA tools, such tools lack for snATAC libraries. *Amulet* leverages the expected read count distributions to identify doublets, while *ArchR* simulates doublets and compares the open chromatin profile of simulated doublets to other barcodes. *Amulet* has the advantage of capturing both heterotypic and homotypic doublets^27^, while simulation-based methods may miss homotypic doublets. It remains unclear to what extent the different ATAC-based methods are able to capture different types of doublets, and future research should investigate this more in depth using simulated datasets with known ground truths. Such work informs on the specificity and sensitivity of different doublet callers and further guides doublet identification.

In the current study we have assessed several data modalities, including single-cell and single-nucleus RNA/ATAC/CITE as well as snMultiome data and shown that *CellDemux* can consistently deconvolve donors. While not tested, we anticipate that *CellDemux* can be extended to deconvolve non-human samples, on the condition that there is sufficient genetic diversity to cluster cells. Moreover, we have tested *CellDemux* on pools containing 3-6 human donors, but we expect that *CellDemux* is equally applicable to libraries with more multiplexed donors. However, the number of singlets captured in droplet-based single-cell data is limited (∼10.000 expected recovered cells in 10X V3 scRNA-seq assays, 8% doublet rate). Multiplexing across too many donors will limit the number of nuclei assigned to each donor and therefore limit the number of informative variants identified in the data. Using *CellDemux*, we found that we were able to confidently identify donors starting from several hundred cells/nuclei onward, sequenced at 25.000 (RNA) and 20.000 (ATAC) read pairs per cell/nucleus.

The ultimate aim of demultiplexing is the assignment of cells to donors. Other demultiplexing methods have provided ways to match cell clusters between different libraries with shared samples, but this does not reveal the identity of the donor^13,15^. Furthermore, *Vireo* identifies a minimal set of discriminatory variants to assign cells to individuals based on quantitative polymerase chain reaction (qPCR), but this approach does not scale well to large cohorts because of time-intensive and laborious experimental work. *CellDemux* requires single-cell sequencing and reference genotyping data. We opted to compare the inferred genotypes to reference, array-based genotypes because of the relatively low costs and experimental burden. Furthermore, the genetic sequence can be re-used in other analyses (genome-wide association studies, expression quantitative-trait loci mapping, etc.). Scripts to compare the genotypes and assign the donor identity to cells are freely available at github.com/CiiM-Bioinformatics-group/CellDemux.

## DATA AVAILABILITY

Single-cell and single-nuclei sequencing data that was previously published is hosted at the European Genome-Phenome Archive (EGA, https://ega-archive.org) (Table 1). Unpublished data will be made available at the time of publication.

**Table 1.**
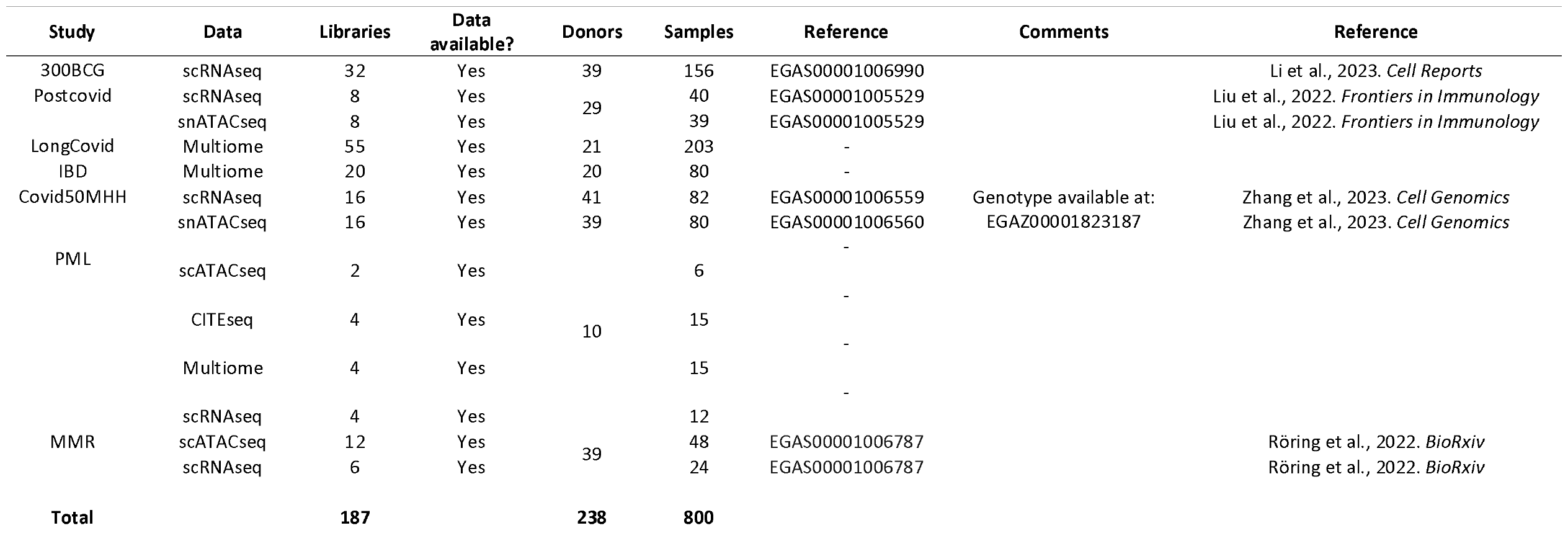
Overview of included datasets in this study.

The *CellDemux* framework is freely available on Github at: https://github.com/CiiM-Bioinformatics-group/CellDemux.git (archived at Zenodo: https://zenodo.org/doi/10.5281/zenodo.10496113) under an MIT open-source license.

## ACKNOWLEDGEMENTS

The authors thank all the participants in the respective single-cell studies.

## REFERENCES

1. Stubbington, M. J. T., Rozenblatt-Rosen, O., Regev, A. & Teichmann, S. A. Single-cell transcriptomics to explore the immune system in health and disease. Science 358, 58–63 (2017).

2. Li, W. et al. A single-cell view on host immune transcriptional response to in vivo BCG-induced trained immunity. Cell Reports 42, 112487 (2023).

3. Schulte-Schrepping, J. et al. Severe COVID-19 Is Marked by a Dysregulated Myeloid Cell Compartment. Cell 182, 1419–1440.e23 (2020).

4. Liu, Z. et al. Multi-Omics Integration Reveals Only Minor Long-Term Molecular and Functional Sequelae in Immune Cells of Individuals Recovered From COVID-19. Front. Immunol. 13, 838132 (2022).

5. Debisarun, P. A. et al. Induction of trained immunity by influenza vaccination - impact on COVID-19. PLoS Pathog 17, e1009928 (2021).

6. Rabold, K. et al. Reprogramming of myeloid cells and their progenitors in patients with non-medullary thyroid carcinoma. Nat Commun 13, 6149 (2022).

7. Zhang, B. et al. Single□cell profiles reveal distinctive immune response in atopic dermatitis in contrast to psoriasis. Allergy 78, 439–453 (2023).

8. Eraslan, G. et al. Single-nucleus cross-tissue molecular reference maps toward understanding disease gene function. Science 376, eabl4290 (2022).

9. Domcke, S. et al. A human cell atlas of fetal chromatin accessibility. Science 370, eaba7612 (2020).

10. Cao, J. et al. A human cell atlas of fetal gene expression. Science 370, eaba7721 (2020).

11. Litviñuková, M. et al. Cells of the adult human heart. Nature 588, 466–472 (2020).

12. Stoeckius, M. et al. Simultaneous epitope and transcriptome measurement in single cells. Nat Methods 14, 865–868 (2017).

13. Heaton, H. et al. Souporcell: robust clustering of single-cell RNA-seq data by genotype without reference genotypes. Nat Methods 17, 615–620 (2020).

14. Kang, H. M. et al. Multiplexed droplet single-cell RNA-sequencing using natural genetic variation. Nat Biotechnol 36, 89–94 (2018).

15. Huang, Y., McCarthy, D. J. & Stegle, O. Vireo: Bayesian demultiplexing of pooled single-cell RNA-seq data without genotype reference. Genome Biol 20, 273 (2019).

16. Lin, X. et al. mitoSplitter: A mitochondrial variants-based method for efficient demultiplexing of pooled single-cell RNA-seq. Proc. Natl. Acad. Sci. U.S.A. 120, e2307722120 (2023).

17. Xu, J. et al. Genotype-free demultiplexing of pooled single-cell RNA-seq. Genome Biol 20, 290 (2019).

18. Stephenson, E. et al. Single-cell multi-omics analysis of the immune response in COVID-19. Nat Med 27, 904–916 (2021).

19. Grant, R. A. et al. Circuits between infected macrophages and T cells in SARS-CoV-2 pneumonia. Nature 590, 635–641 (2021).

20. Yoshida, M. et al. Local and systemic responses to SARS-CoV-2 infection in children and adults. Nature 602, 321–327 (2022).

21. Zhang, B. et al. Altered and allele-specific open chromatin landscape reveals epigenetic and genetic regulators of innate immunity in COVID-19. Cell Genomics 3, 100232 (2023).

22. Köster, J. & Rahmann, S. Snakemake—a scalable bioinformatics workflow engine. Bioinformatics 28, 2520–2522 (2012).

23. Lun, A. T. L. et al. EmptyDrops: distinguishing cells from empty droplets in droplet-based single-cell RNA sequencing data. Genome Biol 20, 63 (2019).

24. Fleming, S. J. et al. Unsupervised removal of systematic background noise from droplet-based single-cell experiments using CellBender. Nat Methods 20, 1323–1335 (2023).

25. McGinnis, C. S., Murrow, L. M. & Gartner, Z. J. DoubletFinder: Doublet Detection in Single-Cell RNA Sequencing Data Using Artificial Nearest Neighbors. Cell Systems 8, 329–337.e4 (2019).

26. Wolock, S. L., Lopez, R. & Klein, A. M. Scrublet: Computational Identification of Cell Doublets in Single-Cell Transcriptomic Data. Cell Systems 8, 281–291.e9 (2019).

27. Thibodeau, A. et al. AMULET: a novel read count-based method for effective multiplet detection from single nucleus ATAC-seq data. Genome Biol 22, 252 (2021).

28. Granja, J. M. et al. ArchR is a scalable software package for integrative single-cell chromatin accessibility analysis. Nature genetics 53, 403–411 (2021).

29. Neavin, D. et al. Demuxafy: Improvement in droplet assignment by integrating multiple single-cell demultiplexing and doublet detection methods. (2022) doi:10.1101/2022.03.07.483367.

30. Curion, F. et al. hadge: a comprehensive pipeline for donor deconvolution in single cell. BioRxiv (2023) doi:10.1101/2023.07.23.550061.

31. Young, M. D. & Behjati, S. SoupX removes ambient RNA contamination from droplet-based single-cell RNA sequencing data. GigaScience 9, giaa151 (2020).

32. Li, H. Minimap2: pairwise alignment for nucleotide sequences. Bioinformatics 34, 3094–3100 (2018).

33. Li, H. Aligning sequence reads, clone sequences and assembly contigs with BWA-MEM. (2013) doi:10.48550/ARXIV.1303.3997.

34. Garrison, E. & Marth, G. Haplotype-based variant detection from short-read sequencing. (2012) doi:10.48550/ARXIV.1207.3907.

35. Dou, J. et al. Single-nucleotide variant calling in single-cell sequencing data with Monopogen. Nat Biotechnol (2023) doi:10.1038/s41587-023-01873-x.

36. Grubbs, F. E. Sample Criteria for Testing Outlying Observations. Ann. Math. Statist. 21, 27–58 (1950).

