## Supplemental Information for "*CellDemux:* coherent genetic demultiplexing in single-cell and single-nuclei experiments"

Supplemental Figure 1

A

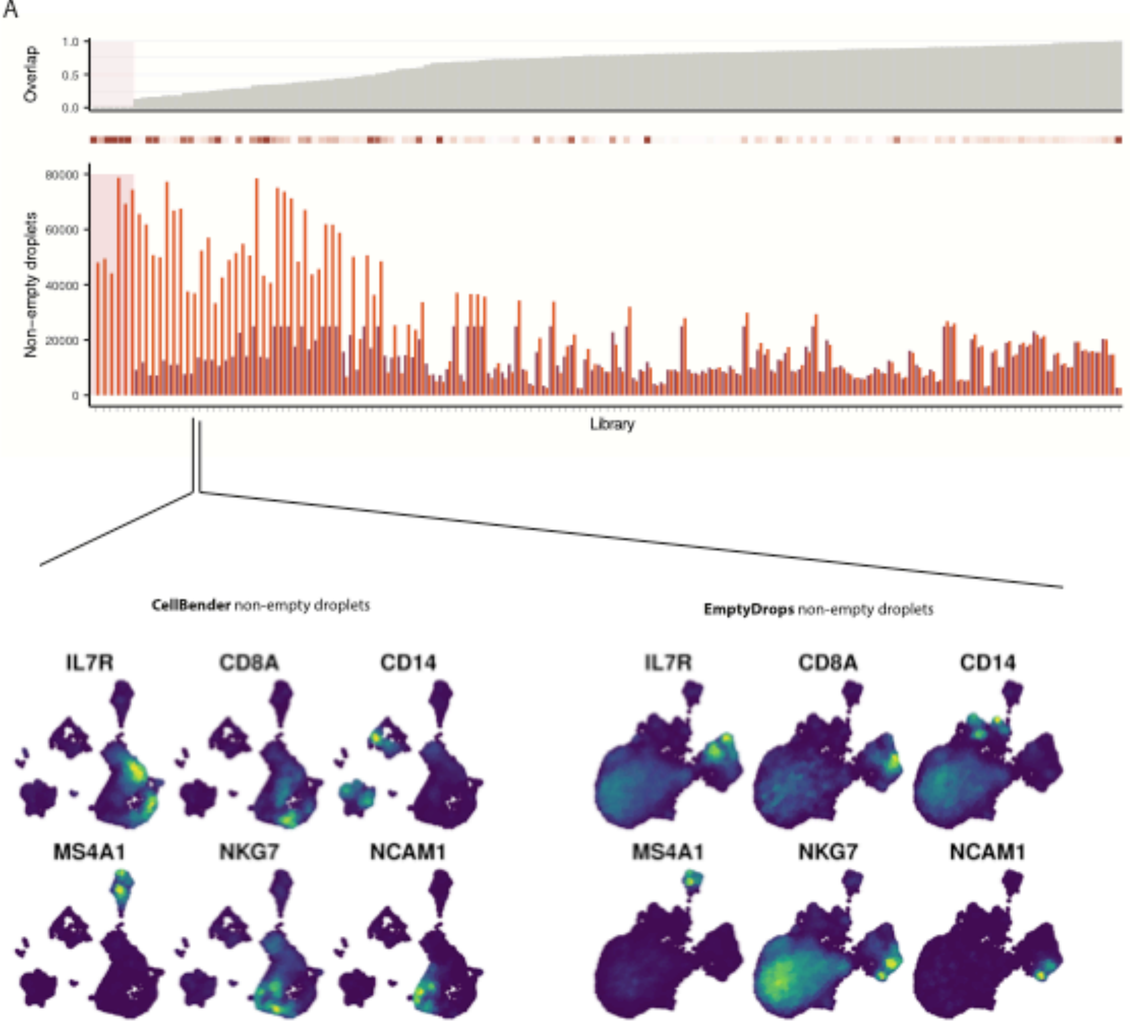


B

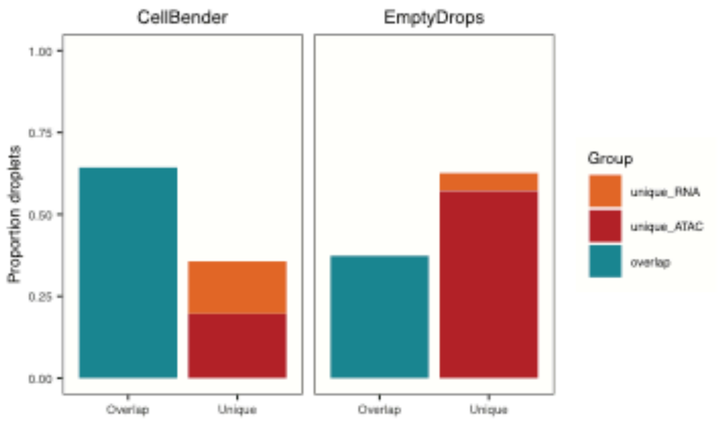

**Supplemental Figure 1. (A)** Example of non-empty oil droplets identified by CellBender and EmptyDrops for one library. The non-empty oil droplets identified by CellBender show more clustering and separation of cell types as identified by marker genes compared to EmptyDrops. **(B)** Testing the identification of non-empty droplets in ATAC data using RNA-based tools CellBender and EmptyDrops. Across 79 Multiome libraries, we show the proportions of droplets identified by either tool to be unique to ATAC, unique to RNA or overlapping between the modalities.

Supplemental Figure 2

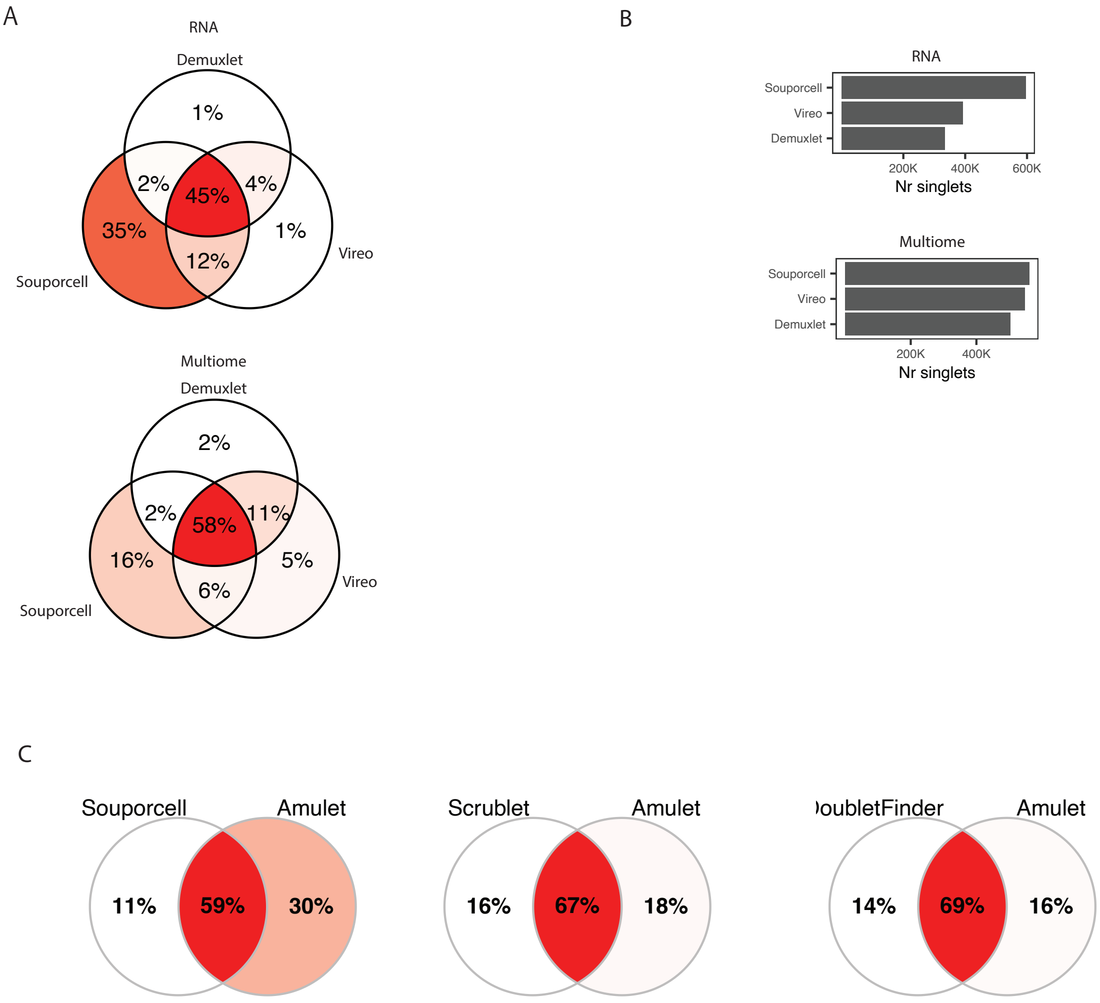


**Supplemental Figure 2. (A)** Venn diagrams comparing the assignment of cell barcodes to donors aggregated across 70 RNA libraries (top) or 79 Multiome libraries (bottom). Barcodes were considered overlapping if they were assigned to the same donor within the same library. **(B)** Total number of singlets assigned to a donor, aggregated across 70 RNA libraries (top) and 79 Multiome libraries (bottom). **(C)** Overlapping singlets between RNA and ATAC as identified by Souporcell/Scrublet/DoubletFinder compared to Amulet.

Supplemental Figure 3

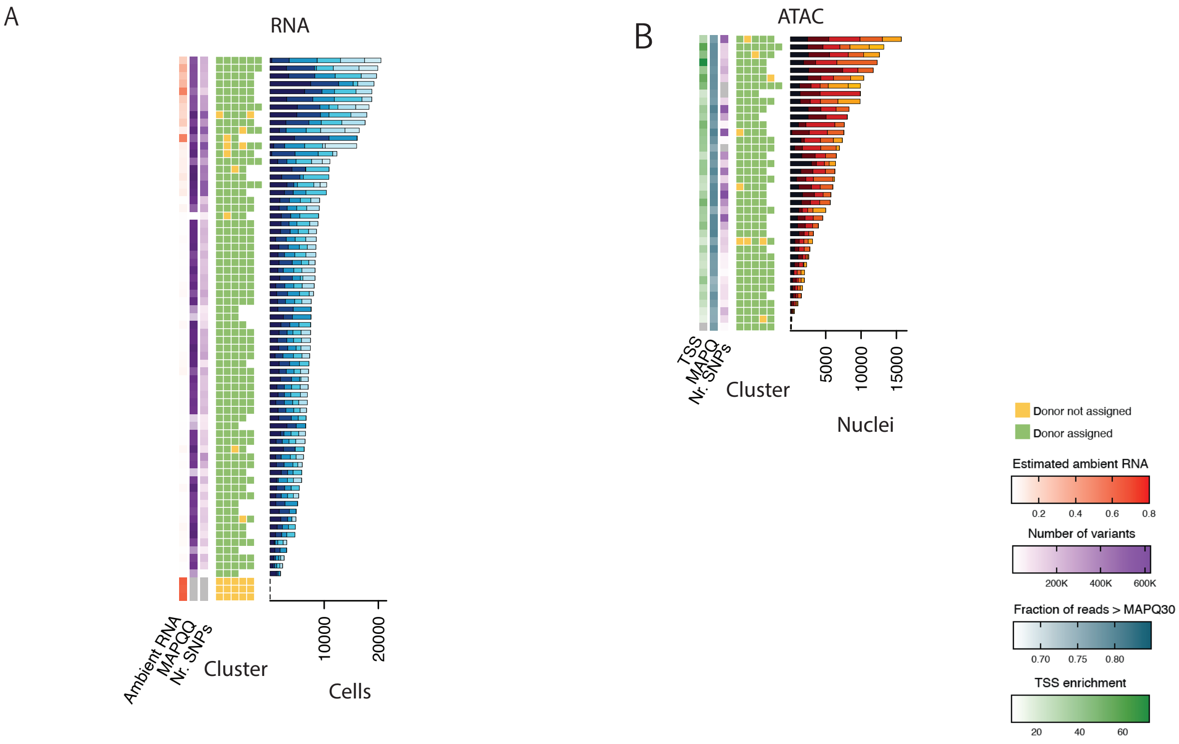


**Supplemental Figure 3.** Heatmaps showing the demultiplexing results for 70 RNA libraries **(A)** and 40 ATAC libraries **(B)**. Each row indicates a library. The main heatmap shows whether a cell cluster is assigned to a donor (green) or not (yellow). We annotate per library several quality control metrics on the left side of the heatmap, the barplot on the right side of the heatmap shows the number of cells/nuclei per cluster.

Supplemental Figure 4

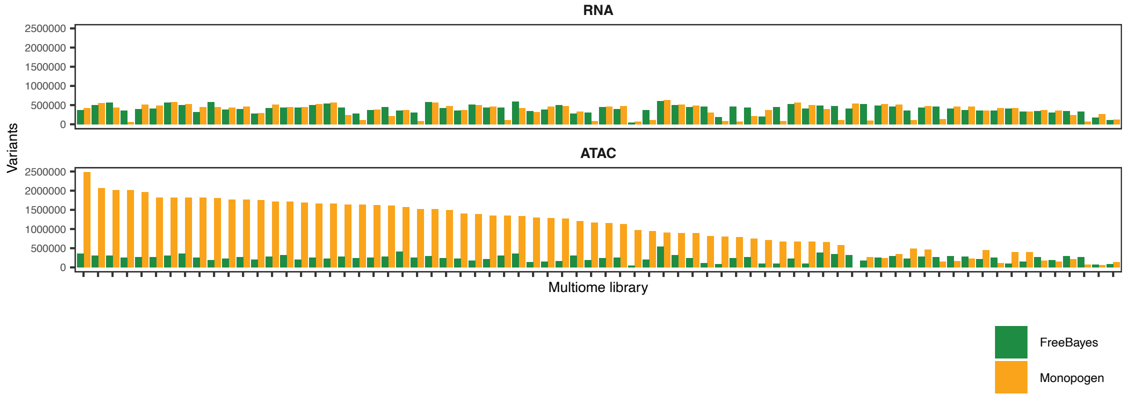


**Supplemental Figure 4**. Number of variants called by FreeBayes (green) and Monopogen (orange) across Multiome libraries. We called variants per library, for RNA-seq (top) and ATAC-seq (bottom) separately.
